# Folding and Persistence Time of Intramolecular G-Quadruplexes Transiently Embedded in a DNA duplex

**DOI:** 10.1101/2021.01.04.425278

**Authors:** Phong Lan Thao Tran, Martin Rieu, Samar Hodeib, Alexandra Joubert, Jimmy Ouellet, Patrizia Alberti, Anthony Bugaut, Jean-François Allemand, Jean-Baptiste Boulé, Vincent Croquette

**Affiliations:** Genome Structure and Instability unit, National Museum of Natural History, Sorbonne University, INSERM, CNRS, 43 rue Cuvier, 75005 Paris; Laboratoire de physique de L’École normale supérieure de Paris, CNRS, ENS, Université PSL, Sorbonne Université, Université de Paris, 75005 Paris, France; Institut de Biologie de l’Ecole Normale Supérieure (IBENS), Ecole normale supérieure, CNRS, INSERM, Université PSL, Paris, France; Depixus, 3-5 Impasse Reille, 75014 Paris; ESPCI Paris, PSL University, 10 rue Vauquelin, 75005 Paris, France

## Abstract

G-quadruplex (G4) DNA structures have emerged as important regulatory elements during DNA replication, transcription or repair. While many *in-vitro* studies have focused on the kinetics of G4 formation within DNA single-strands, G4 are found *in-vivo* in double-stranded DNA regions, where their formation is challenged by pairing between the two complementary strands. Since the energy of hybridization of Watson-Crick structures dominates the energy of G4 folding, this competition should play a critical role on the persistence of G4 *in vivo*. To address this issue, we designed a single molecule assay allowing measuring G4 folding and persistence while the structure is periodically challenged by the complementary strand. We quantified both the folding rate and the persistence time of biologically relevant G4 structures and showed that the dynamics of G4 formation depends strongly on the genomic location. G4 are found much more stable in promoter regions and replication origins than in telomeric regions. In addition, we characterized how G4 dynamics was affected by G4 ligands and showed that both folding rate and persistence increased. Our assay opens new perspectives for the measurement of G4 dynamics, which is critical to understand their role in genetic regulation.

## INTRODUCTION

Nucleic acid sequences with four or more stretches of multiple guanines (GnNxGnNyGnNzGn; n ≥ 2) can form non-canonical four-stranded structures called G-quadruplexes (G4s). These structures are assembled by stacking of coplanar guanine’s quartets (G-quartets) that are stabilized by monovalent cations, such as K^+^ or Na^+^. G4 structures are well described at the structural level. Depending on their sequence, they adopt different folding patterns and can form from the same (intramolecular) or distinct (intermolecular) nucleic acid strands (1–3). Potential G4-forming motifs are widespread across the genomes of a large spectrum of organisms from bacteria to mammals (4–8), and the formation of stable G4 structures *in vivo* has been linked to replication (9–11), oncogene expression (12–14), translation (15–17), DNA repair (18, 19) telomere maintenance (20) or epigenetic marking of a DNA locus (21–23). Questions around the physiological conditions for their formation and their biological roles have made G4 structures the focus of extensive *in vitro* and *in vivo* studies over the last two decades.

Single-stranded (ss) nucleic acid sequences, such as the telomeric 3’-overhang or RNAs, are prone to G4 formation with no challenge from a complementary strand. However, G4 forming sequences are allegedly also found in double-stranded DNA regions, such as promoters (24), minisatellites (25) or replication origins (10). These structures, if stable and unprocessed by helicases (26), may hinder the replication fork and transcription bubble progression, causing fork pausing or promoting DNA breakage (25, 27, 28). Despite causing potential roadblock for molecular motors, current wisdom is that their formation has been harnessed by evolution to encode structural information within DNA, at the expense of evolving protein motors able to remove the stable structures challenging replication (26, 29). Their potential biological effect would require their persistence post replication in a context where they are embedded in a double-stranded DNA (dsDNA) bubble. Persistence of G4 structures within dsDNA is largely unknown, but may depend on their sequence, the genomic context and/or the presence of a bound protein (30–36). Convincing evidences of G4 formation *in vivo* have indeed been reported near several transcription start sites (37) or near-replication origins where G4 structures could participate in transient fork pausing or loading of proteins (10, 11, 38). All these roles suggest that once formed these structures have a significant persistence time in a dsDNA context.

To date, our knowledge of the conditions of G4 formation is largely inferred from *in vitro* biophysical thermodynamic studies of different models G4-forming single-stranded sequences. These approaches have contributed largely to our understanding of G4 stability and structural diversity, but are limited in providing a complete biologically relevant picture of the dynamics of G4 structures at the molecular level. Folding/unfolding of several G4s (such as human telomeric or cMYC promoter sequences) have been studied in recent years using magnetic or optical tweezers (39–43). Most of these studies address the dynamics of G4 in a single-stranded DNA (ssDNA) context. In a non-replicative dsDNA context, the presence of a complementary strand causes a competition with the G4 and the duplex structure. At the thermodynamic level, the energy of hybridization of a Watson-Crick duplex containing a G-rich DNA sequence is much larger (∼50 kcal/mol) than the energy of G4 folding (∼4-8 kcal/mol) (1). Therefore, the competition with a complementarity strand should play a major role in the formation and persistence of G4 structures *in vivo*, as recently pinpointed by Tigran & al (35). Two reports using single-molecule FRET or optical tweezers have shown that G4 can readily compete with the reannealing of dsDNA *in vitro* (44, 45). However, the persistence time of the G4 structure embedded in duplex DNA remains largely unknown.

Here, we developed an original single-molecule setup using magnetic tweezers in order to gain insights into this aspect of G4 dynamics. This setup allows measuring formation and persistence of G4 structures in an alternating context between ssDNA and dsDNA, by opening and closing cyclically single molecule dsDNA hairpins. This label-free assay allowed us to measure the folding time (T_f_) of G4 structures as well as their persistence time (*T*_*p*_) under a defined force cycle. Using this assay, we compared four well-studied model G4 sequences from different genomic regions, namely human telomeric, human promoters cMYC and cKIT, and a replication origin G4 from the chicken genome. Our experiments show that these G4 sequences form structures as expected but show great variation in their folding and persistence times in ways that are not fully described while considering thermodynamic parameters from bulk experiments. Finally, this system also allowed us to assess the effect of a chemical G4 ligand (360B) (46) and an anti-G4 single-chain antibody (BG4) (47) on G4 dynamics. We observed that both molecules favor G4 structure formation by reducing the apparent folding time and increasing the persistence time of the G4. This observation highlights the basic but important idea that some G4s revealed experimentally by such G4 binders may not form significantly in absence of a ligand.

## MATERIAL AND METHODS

### Oligonucleotides and G4 binders

The oligonucleotides used in this study are described in **Supplementary Table 1** and **2**. All oligonucleotides were purchased from Eurogentec (Seraing, Belgium). Lyophilized oligonucleotides were resuspended in distilled water and store at −20°C. The G4 ligand 360B was synthesized in the laboratory by Patrick Mailliet. 360B is similar to the G4 ligand 360A, but the iodine counter ions in 360A are replaced by sulphonates (46, 48) (**Supplementary Data 8**). This modification improves solubility of the compound in aqueous buffers, without affecting its specificity and affinity towards G-quadruplexes (Patrick Mailliet and Jean-François Riou, personal communication). The BG4 single chain antibody (49) was purified and provided by the laboratory of Prof Kevin D. Raney.

### Experimental setup

The PicoTwist™ magnetic tweezers instrument used to manipulate individual DNA hairpins tethered between magnetic beads and a coverslip was described previously (50). Briefly, the magnetic bead was held by a force due to a vertical magnetic field gradient generated by a pair of permanent magnets. Controlling the distance of the magnets from the sample surface allows applying a precisely calibrated variable force on the samples. The movement of these magnets was achieved with the help of PI DC-motor, which allowed us to exert a force with sub-picoNewton accuracy. The force was calculated from the Brownian fluctuations of the tethered bead (51). The DNA substrate used in the single-molecule studies consisted of a 1.1 kb hairpin containing the G4 forming sequence at position 589 from the 5’ end of the molecule (**Supplementary Data 1, 2)**. The hairpin also contains an eight-nucleotide-loop, a 5’-tri-biotinylated ssDNA tail and a 3’ end with 44-nucleotide-long ssDNA tail. We attached the hairpin substrate at 5’ end to a streptavidin-coated magnetic bead and at 3’ end to a glass cover slip through annealing with ssDNA oligonucleotides covalently attached on the surface by click-chemistry. Relevant oligonucleotides for hairpin synthesis are described in **Supplementary Table 1**. To image the beads, a CMOS camera at the image plane of the objective was used to track the position of the magnetic beads in three dimensions with nanometer resolution at 16 Hz. Thus, tracking the z-axis fluctuations of the tethered beads enabled us to monitor the change in extension of the tethered DNA hairpin in real time with an accuracy of about 5 nm. The extension of the DNA changed in accordance with the applied force or by the presence of hybridized oligonucleotides or a G4 structure. Tens of beads were tracked simultaneously in real time in order to obtain statistically significant measurements of single molecule events.

### Experimental conditions for single molecule G4 structure manipulation

All experiments were performed at 25°C in 10 mM Tris-HCl (pH 7.5), with 100 mM KCl or 100 mM LiCl, as indicated in the text and figure legends. Potassium ions (K^+^) strongly stabilize G4s by coordination with the carbonyl oxygens of guanines in the central cavity. To confirm that the detected structures were G4s, control experiments were systematically carried out in 100 mM LiCl. Lithium ion (Li^+^) is too small to fit in the G4 central channel and, hence, stabilizes G4 to a much lesser extent than potassium (52). The experimental system was based on manipulating multiple dsDNA hairpins tethered to a glass slide on one end and to magnetic beads on the other end. Time resolved experiments were started with injection of 10 nM of a 7-base-long oligonucleotide (“blocking oligonucleotide”) complementary to the loop of the hairpin. In experiments using G4 ligands, these were injected concomitantly with the blocking oligonucleotide.

### Data collection and analysis

The image of the bead displayed diffraction rings that were used to estimate its three-dimensional position as previously described (51). From position fluctuations of the bead, both the mean elongation of the molecule and the force applied to it could be deduced (53). The z-axis fluctuations were acquired at 16 Hz. Typical experimental acquisition spanned almost 24 hours, corresponding to ∼2000 open/close cycles (**Supplementary Data 3**).

The experimental mean folding time 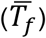 of each G4 sequence was inferred by dividing the whole time spent in the unfolded state (the cumulative time of oligonucleotide blockage before a structure is observed), summed over all beads and over the entire acquisition, by the number of times a G4 structure formed. This average is the best estimator for the parameter of a single exponential law if one considers G4 folding and unfolding as Poisson processes (see **Supplementary Data 4** for explanations on calculations of the parameters and dependency on experiment time). The mean unfolding time, or persistence time 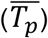, was similarly obtained by dividing the whole time spent in the folded state (> T_hold_ = 15 s) by the number of times a G4 is unfolded (corresponding to the absence of blockage at 0.41 µm and 0.8 µm during a full-force cycle (**Figure1**).

The relative error made on these estimations is 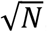, where N is the number of observed events (respectively unfolding and folding events). All results presented below were computed from at least 100 independent G4 events. When two successive cycles showed a blockage at the position of the G4, we considered that it was due to the same structure and that no unfolding and refolding took place during one single hairpin opening phase. If the typical 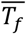 was much larger than *T*_*c*_ (cycle time), the duration of one cycle, the probability of a new G-quadruplex to refold during the opening phase was so low that this assumption does not bias the estimation of 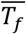. However, in the case where 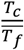 was not negligible, the 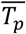 of the G4 could be slightly overestimated as two successive structures could be mistaken for the same structure. It is possible to show (see **Supplementary Data 5**) that the extent of the relative overestimation of 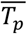 is less than 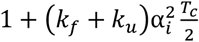, where 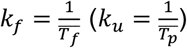 is the real folding (unfolding) rate of the G4 structure and α_i_ = 0.83. This correcting factor was considered in the estimate of the errors, which are for this reason asymmetric. In the same manner, the probability that a G4 structure was formed but could not be detected was considered in the errors relative to its folding time 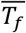.

## RESULTS

### A single molecule assay to visualize G4 formation in real-time

To decipher G4 folding/unfolding dynamics at the molecular level, in a context where G4 structure exists in competition with a dsDNA structure, we developed a single molecule assay based on a magnetic tweezer setup (see Material and Methods and (54)). The principle of the assay used to monitor G4 structure formation is presented in **Figure 1** using data obtained with a G4 sequence from the human cKIT gene promoter (cKIT2) (55). The assay consisted of applying periodic force cycles of 1 minute long which can be separated in three different phases detailed below (**Figure 1A**).

**Figure 1.**
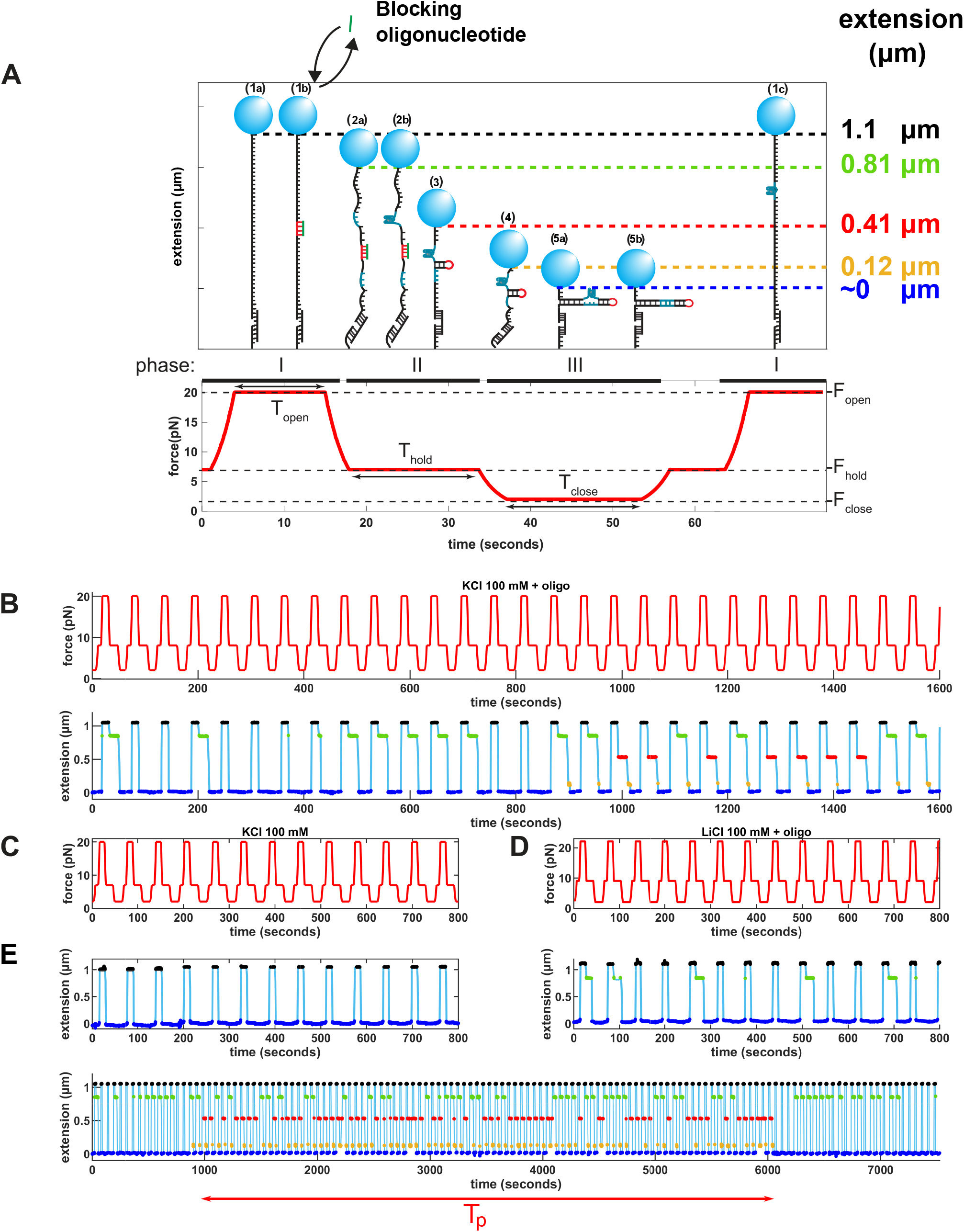
Single molecule assay for measurement of G4 formation. **A**. Description of the force cycle repeatedly applied to each bead and the five detectable extensions. (a), (b) and (c) subscripts denote multiple structures with same extension and thus undistinguishable. **B**. Force cycle profile (red curve) and recorded extension (blue curve) of a single molecule containing cKIT-2 G4 forming sequence in 100 mM KCl plus 10 nM blocking oligonucleotide. The same color code as in A is used to highlight the different extension states. **C**. Same as B in 100 mM KCl without blocking oligonucleotide. **D**. Same as B in 100 mM LiCl with blocking oligonucleotide. **E**. Larger view of D showing a G4 structure folding and unfolding between t=1000s and t=6000s. The persistence time measured by our algorithm for this particular G4 is represented by the underlying red arrow.

#### Phase I: opening the DNA molecule (F_open_ = 20 pN, t_open_=11s)

In order to allow the formation of a G4, the dsDNA must first go through a transient ssDNA state. At 20 pN, the Watson-Crick interactions between the bases are broken and the hairpin unzips into a 1.1 kb ssDNA molecule. This state was characterized by a molecular extension of 1.1 µm corresponding to the full extension of the DNA (**Figure 1A**, structure 1a). At the beginning of the experiment, 10 nM blocking oligonucleotide complementary to the loop region of the hairpin was optionally added to the solution. This amount of blocking oligonucleotide was necessary to delay rehybridization of the hairpin in about every 3 to 4 cycles when lowering the force in Phase II.

#### Phase II: folding the hairpin to evidence G4 structure formation (F_hold_=7 pN, t_hold_=16s)

At F_hold_, depending on the experimental conditions, three extension states may be observed (**Figure 1B**), namely 0, 0.41 µm and 0.81 µm. The 0 µm extension state is the only state observed in the absence of the blocking oligonucleotide (**Figure 1C**) and was attributed to complete folding of the hairpin (**Figure 1A**, structure 5b). In presence of LiCl and a blocking oligonucleotide, we observed an additional extension state at 0.81 µm (**Figure 1D**). This extension was attributed to an unfolded hairpin with the blocking oligonucleotide hybridized to the loop. After a random time distributed within 10-15s, the blocking oligonucleotide spontaneously unbinds and the hairpin can hence refold. In this situation, the G4 may or may not be formed (**Figure 1A**, structure 2a and 2b). The 0.41 µm extension state is observed only in experimental conditions where KCl and a blocking oligonucleotide are present (**Figure 1B** and **1E**). These conditions are favorable for a G4 structure to fold. We interpreted this state as a hairpin blocked from rezipping by the presence of the G4 structure (**Figure 1A**, structure 3). Indeed, a 0.41 µm extension is consistent with the position of the G4 motif within the sequence, it is not observed in presence of LiCl plus blocking oligonucleotide, and it was never observed with a mutated cKIT2 sequence that did not contain a G4 motif (**Supplementary Table 2**).

#### Phase III: lowering the force to 2pN (F_close_) allows G4 encirclement within a dsDNA bubble (t_close_=11s)

Upon further lowering the force, a 0.12 µm extension state may be observed in KCl plus oligonucleotide; we ascribed this extension state to a relaxed state of structure 3 (**Figure 1A**, structure 4). In KCl with blocking oligonucleotide (**Figure 1B** and **1E**), the “0 µm” extension state following a G4 detection (0.41 µm or 0.12 µm states), may correspond, in principle, to a structure where the G4 is embedded within a dsDNA bubble (**Figure 1A**, structure 5a) or where the hairpin is completely refolded (**Figure 1A**, structure 5b). Evidence of the persistence of the G4 within a dsDNA bubble is provided in the next force cycles: if the G4 keeps folded in the hairpin, then rezipping of the hairpin is again blocked at a 0.41 µm extension during phase II. Unfolding of the G4 is therefore simply detected as the first cycle without blockage at 0.81 µm, 0.41 µm or 0.12 µm in phase II. In the recording presented in **Figure 1E**, a single G4 formation event is recorded that lasted for ∼5000 seconds.

### Folding of G4 structures from human telomeres, human oncogene promoters and an avian replication origin

In order to generalize our approach, we tested the behavior of other G4 forming sequences, namely human telomeric repeats (hTelo 21-TTA) (56), the promoter of the human oncogene cMYC (cMYC-Pu27) (57), and a replication origin found in the chicken genome (ori β^A^) (10). **Figure 2A** shows examples of extension traces obtained with hTelo 21-TTA, cMYC-Pu27 and β^A^ origin sequences, in presence of 100 mM KCl and blocking oligonucleotides (corresponding traces obtained during the full course of an experiment from one bead can be found in **Supplementary Data 3**). Formation of G4 structures could be observed for the three different sequences on most tested beads, demonstrating the generality of the assay to study formation of intramolecular G4s. As described above for the cKIT2 structure, experiments were also performed in control conditions (100 mM LiCl with or without blocking oligonucleotide, or 100 mM KCl without blocking oligonucleotide). No blockage corresponding to G4 structures was observed in these conditions for hTelo 21-TTA. We could, however, detect formation of G4 structures, while less frequent and short-lived, in 100 mM LiCl plus blocking oligonucleotide for cMYC-Pu27 and β^A^ origin sequences. Interestingly, the β^A^ origin sequence also exhibited G4 formation in 100 mM KCl without blocking oligonucleotide, but no G4 blockage was observed in presence of LiCl only (**Figure 2B**). These unexpected results are commented in the next section.

**Figure 2:**
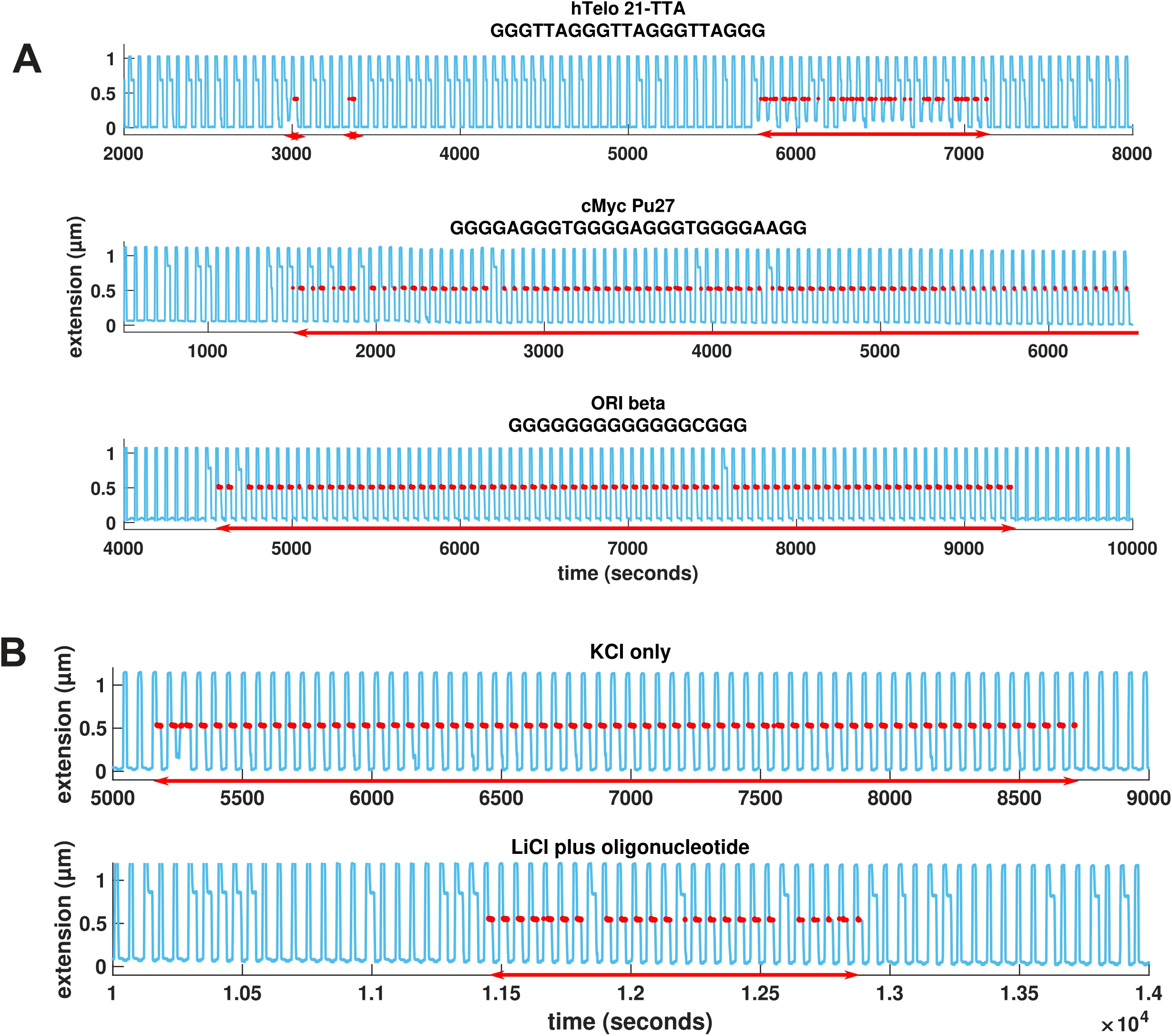
Extension traces from single molecules containing hTelo 21-TTA, cMYC Pu27 or β^A^ origin G4 forming sequences. **A**. hTelo 21-TTA (top), cMYC (middle) and β^A^ origin (bottom) G4 forming sequences in 100 mM KCl and 10 nM blocking oligonucleotide. **B**. β^A^ origin G4 forming sequence in KCl without blocking oligonucleotide (top) and in LiCl plus blocking oligonucleotide (bottom). Formation of a G4 structure is detected by a blockage (red) in phase II, as described in Figure 1. Measured G4 persistence times in the represented traces are highlighted by red arrows.

### Comparative folding and unfolding dynamics of intramolecular G4s

We reasoned that folding time of the structures and their persistence could be inferred from our assay. To gain insights into the folding and unfolding dynamics of the G4 structures, we recorded multiple events of G4 formation over several beads for each G4 forming sequence (more than 100 recordings per sequence, each experiment lasting up to 16 hours). We were able to monitor a field of multiple molecules at the same time to obtain statistically significant measurements of the G4 folding and persistence times under various conditions.

For each sequence studied, we compiled the experimentally folding times (*T*_*f*_) as all the times where the blocking oligonucleotide delayed hairpin refolding in phase II (extension of 0.81 µm at 7 pN) before first observing the formation of a G4 structure (first blockage at 0.41 µm) (**Figure 1A** and **Supplementary Data 4**). The average folding time 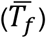 of a given sequence was therefore obtained by dividing the sum of T_f_ over all beads and over the entire acquisition, by the number of times a G4 structure was formed. Thus, the inverse of 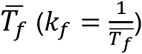 is a direct measure of the probability of folding a G4 structure per unit of time spent at 7 pN. A longer average folding time indicates a smaller folding probability while a shorter average folding time indicates a larger folding probability. Similarly, we determined persistence times (*T*_*p*_) of a G4 as the times spent by a molecule in a G4 conformation before it unfolded (**Supplementary Data 4**). The average persistence time 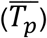 is acquired by dividing the sum of *T*_*p*_ over all beads and over the entire acquisition, by the number of times a G4 is unfolded (corresponding to the first absence of blockage at both 0.81 and 0.41 µm extension following detection of a structure). Therefore the 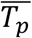 represents the stability of a given G4 structure in a context alternating between ssDNA and dsDNA. The inverse of 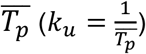 is therefore the unfolding rate of the G4 structure.

Comparison of the dynamic properties of the different G4 structures is shown in **Table 1** and **Figure 3A**. The four studied G4 sequences (hTelo 21-TTA, cKIT2, cMYC-Pu27 and ori β^A^) displayed large difference on their 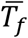 (from ∼30 to ∼12000 seconds) and 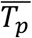 (from ∼500 to ∼11000s) at 25°C in 100 mM KCl plus blocking oligonucleotide. Some G4 sequences, such as cMYC-Pu27 and β^A^ origin, had a very short folding time and very long persistence time (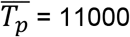 and 9000s respectively). In comparison, cKIT2 exhibited a longer folding time 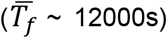 and a shorter persistence time 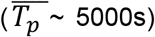 compared to cMYC-Pu27 and ori β^A^. Finally, hTelo 21-TTA presented the most dynamic structures, with relatively short folding 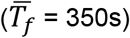 and short persistence times 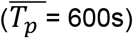.

**Table 1:** Average folding and persistence times of G4 structures in different conditions. * “nd” stands for “not determined”.

| | Name | cMYC - Pu27 | cKIT2 | ori $\beta^A$ | ori $\beta^A$ - m12 | ori $\beta^A$ - m16 | 21-TTA | 45-TTA | 21-CTA |
| --- | --- | --- | --- | --- | --- | --- | --- | --- | --- |
| Average Folding Time ($\overline{T}_f$) | LiCl | nd* | nd | no G4 | nd | nd | nd | nd | nd |
| | LiCl + oligo | 2300 ± 200 | no G4 | 3000 ± 350 | no G4 | no G4 | no G4 | no G4 | no G4 |
|  | KCl | no G4 | no G4 | nd | no G4 | no G4 | no G4 | no G4 | no G4 |
| | KCl + oligo | 90 ± 10 | 12000 ± 1700 | 30 ± 13 | 1400 ± 170 | 9000 ± 1300 | 350 ± 30 | 150 ± 13 | 1700 ± 130 |
| Average Persistence Time ($\overline{T}_p$) | LiCl | nd | nd | no G4 | nd | nd | nd | nd | nd |
| | LiCl + oligo | 400 ± 50 | no G4 | 650 ± 80 | no G4 | no G4 | no G4 | no G4 | no G4 |
| | KCl | no G4 | no G4 | 9000 ± 1200 | no G4 | no G4 | no G4 | no G4 | no G4 |
| | KCl + oligo | 11000 ± 1700 | 5000 ± 900 | 9000 ± 1300 | 6500 ± 1300 | 125 ± 20 | 600 ± 80 | 500 ± 50 | 350 ± 60 |

**Figure 3.**
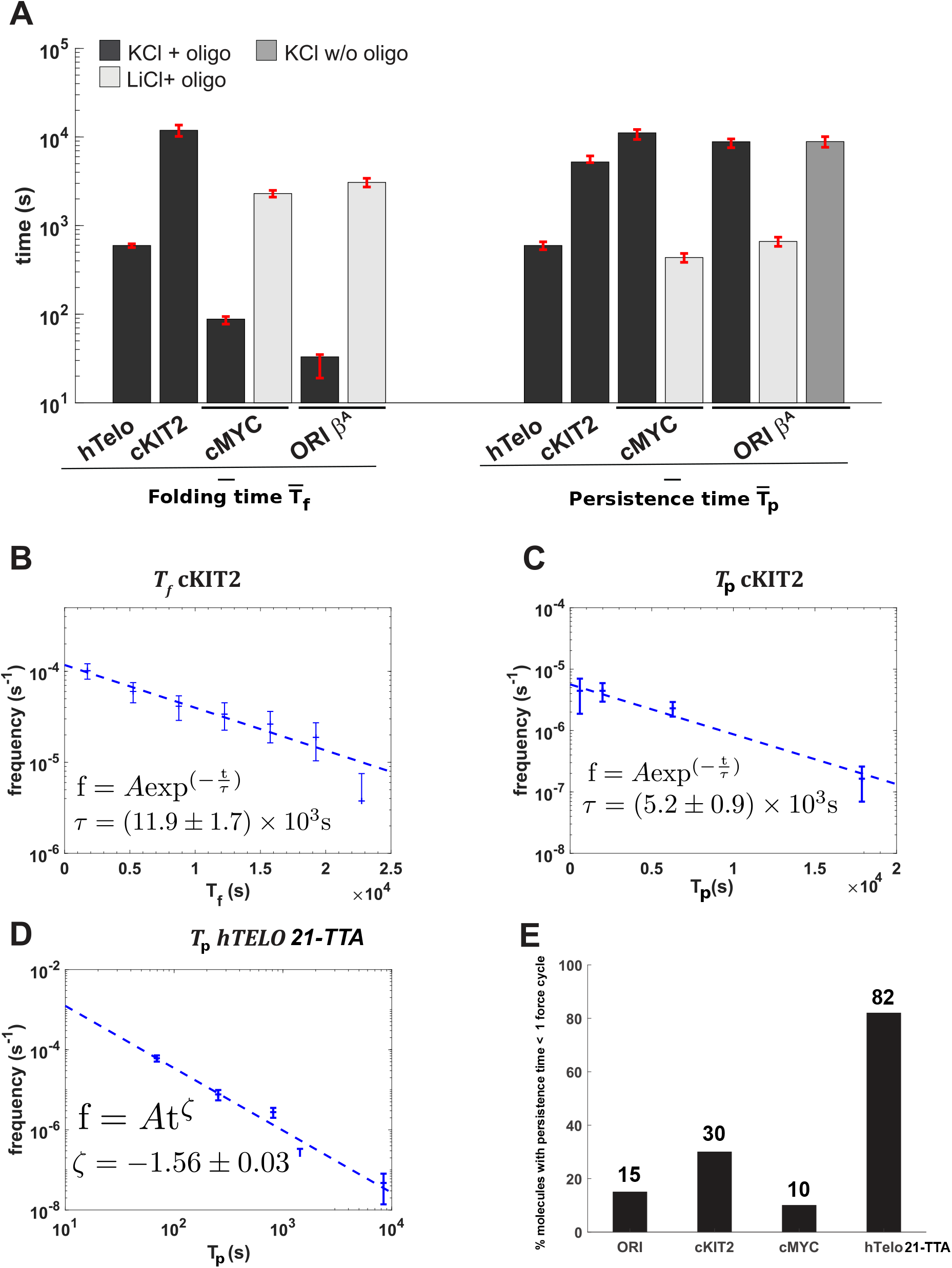
Folding time and persistence time of G4 structures. **A**. Average folding times and persistence times of various G4 structures. Frequency of experimentally measured folding times (**B**), persistence times (**C**) and first order exponential fits for the cKIT2 G4. **D**. Frequency of experimentally measured persistence times of hTelo 21-TTA G4 and power law fit of the data. **E**. Percentage of G4 structures with a persistence time shorter than one force cycle.

The distribution of folding (*T*_*f*_) and persistence (*T*_*p*_) times followed in most of the cases an exponential law, as exemplified for the cKIT2 G4 in **Figure 3B and 3C**. This suggests folding and unfolding of the G4 in a single step. However the persistence time of the hTelo 21-TTA G4 is best fitted by a power law (**Figure 3D**), which would indicate a mixture of indistinguishable conformations with different 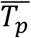, corroborating previous results pointing to structural diversity of human telomeric G4 (43).

By design, one force cycle corresponds to the lower-resolution limit of our assay. Therefore, structures that were stable less than one cycle (≤ 15s) could represent unstable structures either with a short persistence time or structures that are mechanically unfolded during the pulling step at 20 pN. These ambiguous events were thus not considered in the calculation of the 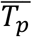. Such unstable structures were frequently observed for the telomeric G4 (21-TTA), since 82% of the structures observed had a persistence time shorter than one cycle. For cMYC-Pu27, cKIT2 and β^A^ origin G4s, the structures lasted less than one cycle represented respectively 10%, 30% and 15% of the events observed (**Figure 3E**). Interestingly, as noted above, we could also observe for the ori β^A^ G4 and the cMYC-Pu27 G4 a blockage at 0.41 µm extension in LiCl in presence of a blocking oligonucleotide. However, the average folding time of the structures was longer (26-fold increase for cMYC-Pu27 and > 90-fold increase for ori β^A^), and their average persistence time was significantly shorter (respectively 26-fold and 13-fold for cMYC-Pu27 and ori-β^A^) than the ones obtained in KCl plus blocking oligonucleotide. The lower persistence of the structures observed in LiCl compared to KCl confirms the involvement of G4 structures in blocking the hairpin refolding. However, we cannot completely rule out the contribution of a structure forming on the opposite C-rich strand, namely a i-motif, although the neutral pH buffer conditions used in our assay should disfavor its formation (57). In the absence of the blocking oligonucleotide and in 100 mM KCl, ori-β^A^ could form a structure with similar persistence (9000s) than the one in KCl plus oligonucleotide. This result suggests two additional possibilities: the β^A^ origin sequence might not require the opening of double-stranded DNA to form a G4 or it may fold in single-stranded context even at high force (∼20 pN).

### Folding time and persistence time of ori β^A^ mutants

Sequences composed of long guanine stretches (such as β^A^ origin sequence) form preferentially intermolecular G-quadruplexes in bulk experiments, rendering difficult to assess the effect of intramolecular G4 folding. Our single-molecule assay helps to overcome this issue. In order to test the sensitivity to mutation of the ori β^A^ sequence in our assay, we measured the dynamics of mutated sequences of ori β^A^ containing an adenine instead of a guanine at position 12 (m12) or at position 16 (m16) (**Table 1** and **Supplementary Table 2**). These point mutations were previously reported to lower the ability of these sequences to form G4 structures and were shown to correlate with lower replication origin activity *in vivo* (10). In our assay, the two ori β^A^ mutants could still fold into a G4 structure, although they exhibited a much longer folding time (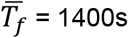 for m12 and ∼9000s for m16), respectively a 42-fold and 270-fold increase compared to wild-type (wt) ori β^A^. The ori β^A^-m12 G4 structure exhibited a long persistence time 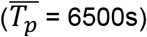, similar to the wt ori β^A^ structure, while β^A^ ori-m16 G4 had a much shorter 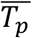 (125s) compared to WT and to m12. In addition, m16 presented a 4.5-fold increase in the percentage of G4 structures with very short persistence time (<1 force cycle) compared to wt and to m12 (69 % for m16 instead of 15 % for wt and m12) (**Supplementary Data 6)**. These results show that our assay can reveal intramolecular G4 dynamics and the effect of subtle sequence difference in G4 forming sequences made of long guanine stretches, in a way not easily predictable by sequence analysis (10). In the case of the ori β^A^ mutants, we see that the guanine at position 12 only affects the folding rate of the G4, whereas the guanine at position 16 is critical for the efficient folding and persistence of the G4 structure in the dsDNA context.

### Folding time and persistence time of dimeric human telomeric repeats and of the common CTAGGG variant

Human telomeres contain variant repeats interspersed with the consensus TTAGGG repeat. The CTAGGG variant, when present as a short contiguous array within the telomere, causes telomere instability in the male germ line and somatic cell (58). It has been shown that four successive CTAGGG repeats were prone to form a G-quadruplex *in vitro*, and adopted a different G4 conformation than the wt (TTAGGG)_4_ repeats in KCl (59). In our assay, the formation of the 21-CTA variant in KCl was five times slower than the wt 21-TTA G4 (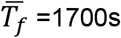 versus 350s). The persistence time of the variant 21-CTA G4 was also reduced by 2-fold compared to the wild type (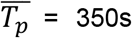 versus 600s) (**Supplementary Data 7B and Table 1**). Therefore, the variant cytosine affects the folding ability of the G4 and has a modest effect on its persistence time. Altogether, these results suggest that the CTAGGG variant has a slightly lower ability to fold into a G4 than the wt telomere repeat. This result does not support the idea that a difference in G4 folding properties would be responsible for the increased genomic instability where this variant is found at human telomeres.

Since telomeres are composed of tandem repeats *in vivo*, we also measured the folding and unfolding dynamics of one versus two G4 units (45-TTA) (**Supplementary Data 7B**). Folding of this sequence into a G4 took in average half the time 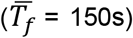 compared to the 21-TTA sequence in KCl. This difference in mean folding time between one and two G4 units is compatible with independent folding of the two G4 structures, suggesting no or little interaction of the tandem G4s in our system. The 45-TTA G4 displayed a persistence time 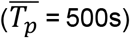 similar to the one of 21-TTA G4 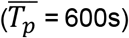, indicating again a lack of a cooperative effect on the persistence time of the structures. The decrease in folding times suggests a higher probability to form the G4 structure in the case of the repeated sequence 45-TTA. This is compatible with the larger number of G4 nucleation sites permitted by the repeat of G4 motifs (**Supplementary Data 7C)**. Among these five nucleation sites, three of them (position 2/3/4) prevent the formation of two consecutive G4 structures. Thus, the probability that the 45-TTA sequence folds into a unique G4 structure is higher than the probability to form into two consecutive ones. However, our assay does not allow distinguishing effectively between the formation of one or two structures in this case.

### G4 stabilizing ligands affect both G4 folding and unfolding dynamics

Finally, we wanted to assess the effect of G4 ligands on the kinetics of G4 folding and persistence in our assay. The potential to manipulate G4 formation at telomeres or oncogene promoters have made them an attractive target for antitumoral drug design (60–63). Small molecules and antibodies have been developed for targeting G4 *in vitro* and have been widely used to evidence G4 formation *in vivo* and potentialize their biological effect (64–66). We tested how such molecules affected folding and unfolding dynamics of G4 structures in the context of our single molecule assay, using the CTA telomere variant as a model sequence (**Table 2**). As demonstrated above, this sequence has the benefit of exhibiting a long folding time 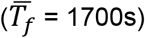 and a short persistence time 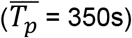, therefore allowing any effect of the ligand on these two values to be easily measurable.

**Table 2:**
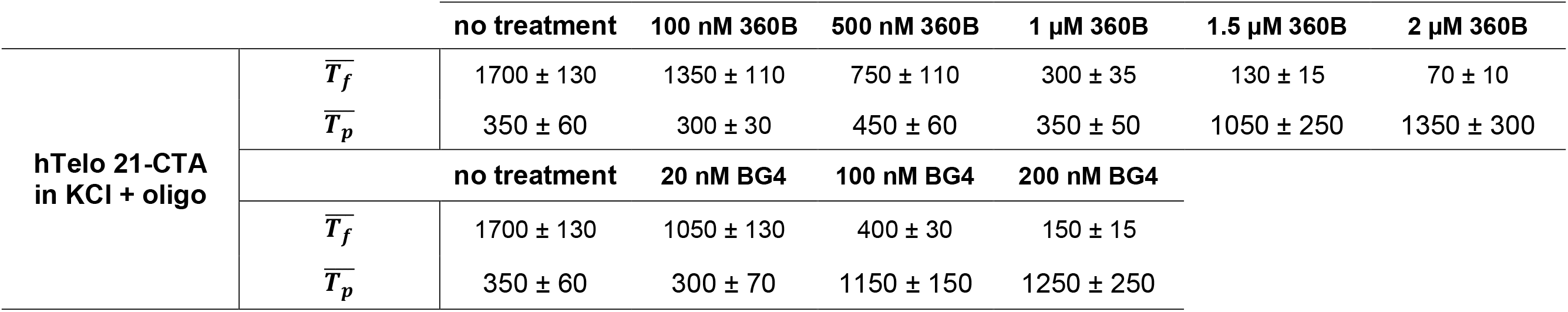
Average folding and persistence times of hTelo 21-CTA G4 structure at different concentrations of 360B ligand and BG4 antibody.

We first investigated the effect of 360B, a selective G4 ligand of the pyridine derivative series, on both folding and unfolding dynamics of the CTA variant G4. The pyridine dicarboxamide derivative 360B is a more water-soluble formulation of the commonly used ligand 360A (**Supplementary Data 8**)(46, 67). We measured the folding and persistence times of the G4 in 100 mM KCl and blocking oligonucleotides with increasing concentrations of 360B. First, we observed that the percentage of structures lasting less than one open/close cycle decreases from 88% to 45% when the ligand concentration increases from 0 to 2 µM, confirming the stabilization of the structures (**Supplementary Data 8A**). Second, increasing concentration of 360B affected both the folding time and the persistence time of the structure in a dose-dependent manner. With increasing concentrations of 360B, we observed folding faster (decreased 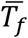) and more persistent G4 structures (increased 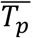), suggesting that the ligand favors the formation of the G4 structure and increases its persistence time in a dsDNA context (**Figure 4A**). Interestingly, 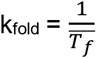 increased non-linearly with the concentration of ligand. The data fit to a quadratic polynomial equation (R^2^=0.9957), which would suggest that two ligand molecules bind simultaneously on one G4 structure. This is reminiscent of what has been reported previously with the interaction between human telomeric G4 and the ligand Cu^II^-tolyterpyridine (68). The ligand also affected the persistence time of the structure. Indeed, 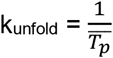 decreased with increasing concentration of 360B, with the data best fitted by a first order exponential decay (**Figure 4C**). We also tested the effect of 360B on the cKIT2 G4, a structure which exhibits long 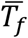 and long 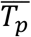. Similarly to the 21-CTA G4, we observed a faster folding time and a longer persistence time of the cKIT2 G4 in presence of the ligand. 100 nM of 360B was sufficient to reduce 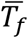 by 5-fold and increase 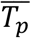 by more than 20-fold. Because of its already long persistence time, most of the cKIT2 G4 (∼60%) were already not unfolded during the recorded time of the experiment (>60,000s) at a concentration of 100 nM of 360B (**Supplementary Data 9**).

**Figure 4:**
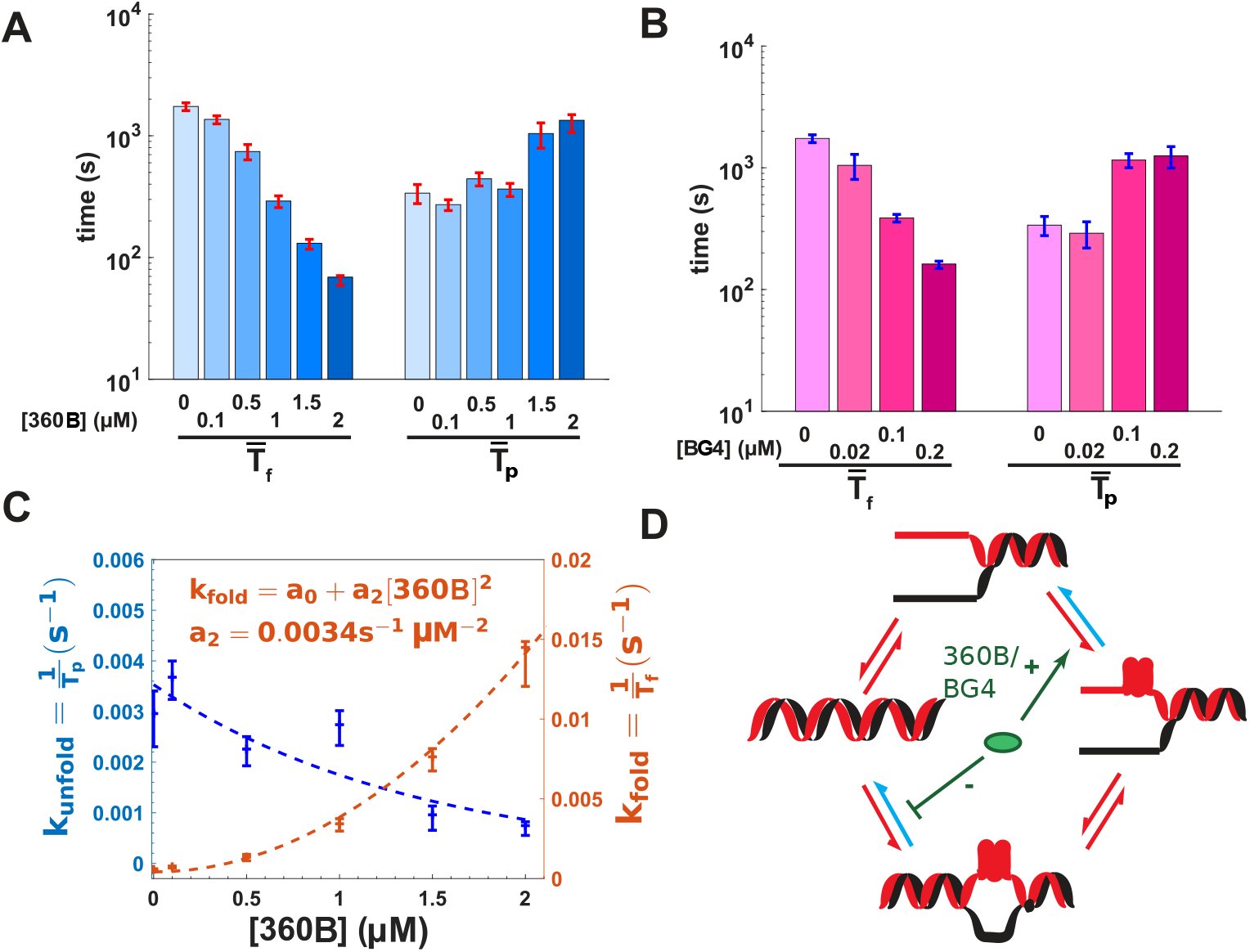
Effect of G4 ligand 360B and BG4 antibody on G4 folding and persistence times. **A**. Folding time and persistence time of 21-CTA telomeric variant G4 with increasing concentration of small ligand 360D. **B**. Folding time and persistence time of 21-CTA telomeric variant G4 with increasing concentration of BG4 antibody. **C**. Kinetic parameters of folding and unfolding of the 21-CTA G4. Fitting equations of the experimental data are displayed. **D**. Kinetic steps of folding a G4 into double-stranded DNA. G4 ligand and antibody (green ellipse) may affect the folding as well as the unfolding kinetics of the G4 structure.

Finally, we tested the effect of the anti-G4 single chain BG4 antibody (65, 69) on the 21-CTA sequence. Similarly to our results with 360B, BG4 elicited a dose-dependent effect on both *T*_*f*_ and *T*_*p*_ (**Figure 4B and Supplementary Data 8B**). Therefore, the two ligands behave phenomenologically in our assay by favoring G4 formation in ssDNA and slowing down its opening, as schematized in **Figure 4D**.

## DISCUSSION

In this work, using a single molecule DNA manipulation assay, we compared the dynamics of G4 folding and unfolding for several G4 forming sequences in a context where single-stranded DNA exists in equilibrium with dsDNA. These results are an important step towards deciphering the relationship between features of G4 forming sequences and their potential to form and persist as a G4 structure in a dsDNA context. Here, we studied well-known G4 motifs belonging to three different genomic loci, including human telomeres, human oncogene promoters of cMYC and cKIT genes, and a chicken replication origin.

Our textbook view of G4 formation in genomic DNA is that the folding of such structure needs a transient ssDNA intermediate, which could likely occur during replication, transcription or DNA repair (70). Indeed, to form in our assay, most structures required a step in a single strand conformation at low force. There was, however, one significant exception with the β^A^ origin G4, which folds into a very stable intramolecular structure even in the absence of a blocking oligonucleotide. Importantly, this is the first time that an intramolecular G4 formed by a long guanine stretches like the β^A^ origin sequence was experimentally observed, since they tend to fold into intermolecular G4 structures (or a mixture of intra- and intermolecular structures) at the DNA concentrations required for common *in vitro* spectroscopic studies (10). This propensity to fold is unique among the sequences tested here and raises the possibility of the contribution of a i-motif forming on the opposite strand to the overall dynamics of the structure evidenced by our assay (35). Although analysis of i-motif formation is beyond the scope of the present study, our assay opens interesting avenues to assess the dynamics of its formation in slightly acidic buffers where its formation would be favored (71). Finally, though our label-free assay is a simplistic model with regard to an actual *in vivo* situation, the observed dynamics of the G4 structures are consistent with their purported role *in vivo*. In the case of a replication origin G4, fast folding and long persistence time in a dsDNA should be advantageous if the structures serve as anchors for protein complexes prior to replication licensing as previously proposed (38). Although not forming as easily as the β^A^ origin G4, the two promoter G4s, cMYC and cKIT, also displayed long persistence in dsDNA, which is consistent with a sustained effect on transcription activity (72).

At the other end of the spectrum, human telomeric sequences were the weakest of the tested structures. They were rapid to form, but showed limited mechanical stability under a 20 pN force and exhibited relatively short persistence time, compared to other sequences. The short persistence (less than one force cycle) observed with human telomeric G4s (**Supplementary Data 7A**) can be attributed either to a low mechanical stability at high force (20 pN) or to a low persistence within a dsDNA bubble at zero force. Although we cannot distinguish between these two possibilities, previous single-molecule assays have shown that the human telomeric G4 repeat could fold into four different G4 conformations, three of them being unstable at 20 pN (43). The distribution of persistence times measured with our assay was best fitted by a power law, also suggesting the contribution of several structures. Our results corroborate other former single molecule studies (43) and point out that a single human telomeric G4 is a dynamic structure, forming promptly in ssDNA but also dissolved rapidly under mechanical forces or in presence of a complementary strand. In the human genome, the telomeric sequence TTAGGG is mainly embedded in a large tandemly repeated dsDNA regions which may form multiple consecutive G4s. This raises the possibility that the impediment to molecular motors when progressing through telomeric repeats owes little to the stability of individual G4 structures alone, at least in the dsDNA portion of telomeres, but probably mainly to other factors, like DNA-bound proteins.

Our results using G4 stabilizing ligands showed that the 360B ligand and the BG4 antibody significantly increased the formation rate of the G4s, as well as their persistence time, proving that they favor G4 formation in addition to stabilizing the structure. This leads to a cautionary remark, when such compounds are used to demonstrate the presence of G4 structures *in vivo*. Indeed, their effect on the folding rate of G4 structures limits their use towards characterizing physiological G4 folding in live cells (73). In this regard, our assay could help characterize ligands or antibodies with limited effect on folding rates, which would be useful for in vivo physiological studies of G4 dynamics.

## Conclusion

Contrarily to previous single-molecule measurements that were performed at constant forces and without the presence of a complementary strand (39–45), our label-free assay allows *in vitro* measurement of G4 folding and persistence times in a context that simulates the competitive environment of G4 structures formation *in vivo*. This assay could be easily adapted to study various biochemical processes (in first place replication and transcription), by tuning the parameters of the force cycles and by studying the effect of proteins involved in G4 processing. It is noteworthy that such modifications in the force cycles significantly affect the kinetics of formation but not the persistence time of a G4 structure (**Supplementary Data 10**). Therefore, our assay allows assessing a wide range of G4 folding dynamics while the DNA molecule is subjected to different mechanical stresses or G4-interacting ligands. Finally, gathering folding kinetics and persistence of a wider array of sequences in this assay could help improve the predictive power of computational methods developed to find G4 forming sequence in genomic data and score their ability to form a stable structure in a dsDNA context (74).

## AVAILABILITY

The data that support the findings of this study are available on request from the corresponding authors.

## SUPPLEMENTARY DATA

In accompanying document

## ACKNOWLEDGEMENT

The authors thank Patrick Mailliet for providing the 360B ligand and Kevin D. Raney for providing us with the BG4 antibody. The authors thank Bertrand Ducos and Jessica Valle Orero for discussions Jean-François Riou and Nicolas Desprat for fruitful comments on the manuscript.

## CONTRIBUTION

PLTT performed the experiments. PLTT and AJ designed and made the hairpin molecules. PLTT, MR, JBB, VC analyzed the data. PLTT, JBB, MR wrote the manuscript, with contribution from PA and AB. SH, VC, JBB, PLTT, MR designed the experiments. JFA and VC constructed the magnetic tweezers. All authors discussed the results and reviewed the manuscript before submission.

