## supplemental data and tables for "Folding and Persistence Time of Intramolecular G-Quadruplexes Transiently Embedded in a DNA duplex"

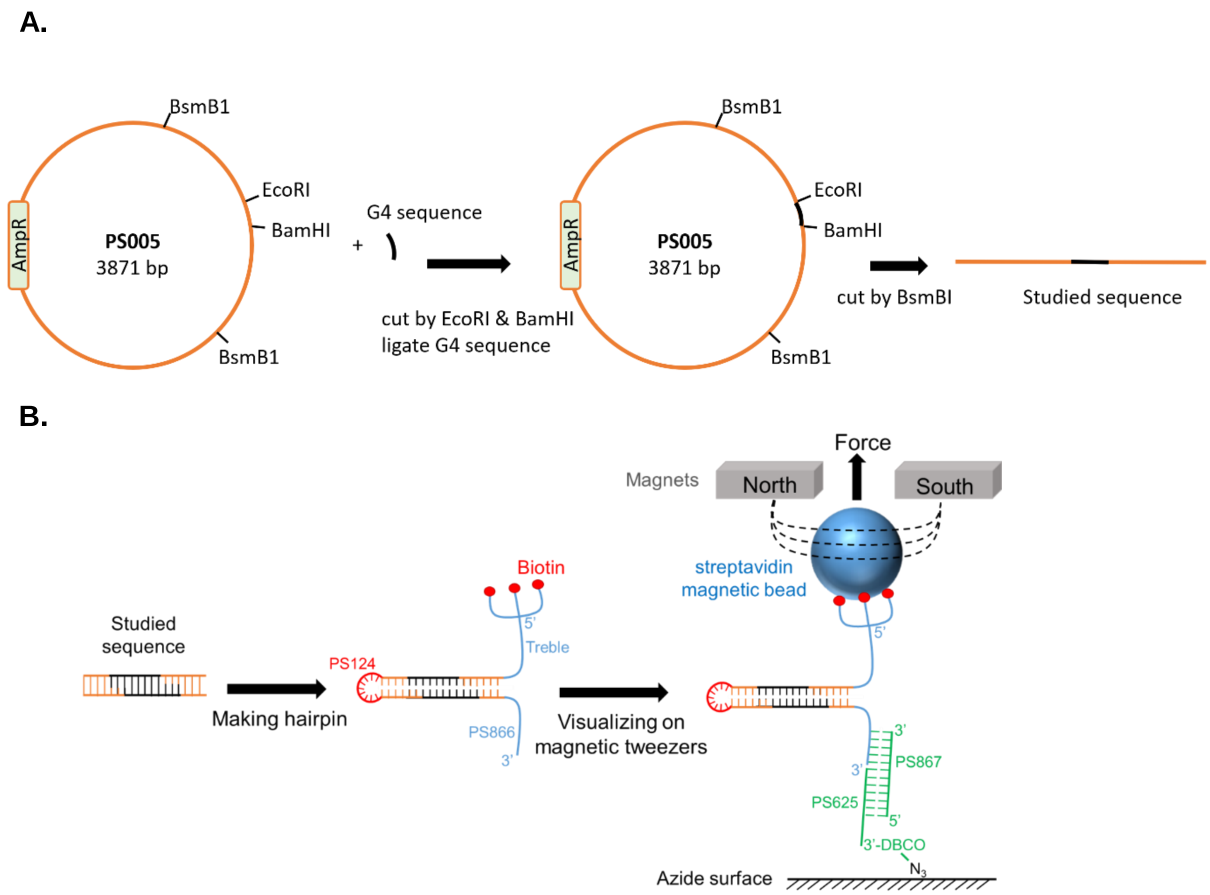


**Supplementary Data 1: Synthesis scheme of the single molecule substrate. A. Cloning of a G4 sequence in the PS005 plasmid.** Oligonucleotides containing respectively the G-rich sequences and their complementary strands were incubated at equimolar ratios at 95°C in 1X Tris-EDTA buffer (pH 7) and digested by EcoRI and BamHI. The PS005 vector was digested by the same restriction sites and ligated with the double-stranded insert containing the potential G4 forming sequence (**Supplementary Table 2**). The recombinant plasmid containing the G4 sequence was then amplified in *E. Coli*. The purified plasmid was then digested with BsmB1, which generates a 1.1 kilobase pair DNA fragment containing the G4 motif and non-palindromic overhangs on each end. **B. Diagram of the hairpin synthesis scheme and sequences involved in tethering the DNA hairpin on azide coated glass coverslip.** The double-stranded DNA fragment was ligated on one end with a loop sequence (PS124) and in the other end with a Y-shape molecule resulting from the annealing of DNA oligonucleotide PS 866 to the treble oligonucleotide (**Supplementary Table 1**). The DNA hairpin contains three biotins at the 5'-end which allow a strong binding on a streptavidin-coated magnetic bead. The DNA molecule is tethered to a glass cover slip through annealing to oligonucleotide PS867 at 25°C, which itself is annealed to oligonucleotide PS625. PS625 is covalently linked at its 3' end to the coverslip through a dibenzocyclooctyne group (DBCO)-azide reaction.

5'BiotinATGGTCAAGCATGCCGCTTTTCGGTTCCCGTGTGTCTTTTGGTCTTTCTGGTGCTCTTCgaatggagAcGAGCTCAGGCCTTAGAGTCAAGACTACAGTAGATACTAATG
ACTGACAGTACCATCAGCATCTCTATATGTAACAGAGTCTTAGAACTGTAGTACTCATGTCATGATCAGATCCATCTTGATGTACTGTCAGACTATCCTCATAAGTCTAGTGAGTAC
TGAAGATACATAGCTCAGATCTGATCATACACGACAGGAGTAGAAGTATACACTGAAGTAGATAGATCGATTGACTATGACCATGCTATGACTGACTGCTACTATTCTAGAGTGA
CAGTATTACTAAGTACGAGCATCTAACAAGATGAACCTGACAGACACTGACATAGCGATTATCATAGCTAGTCACTCTGACGTATCTCAGTCTAACATCAGCACATGAATGTATGA
CTACAGTTACTATCAAGTACACTCTGAGTCAGCATCATCAGTAGATCGATCAGTACAGTATCTAGGACATGACTACTTGAGTGAGAGAATCTTCTACTATCTGTAGTCAGCTTCGT
AACACTCTGATGTCATCTTAATGCTATGATTGTCTACTTGATCGACTACTAGATAACTATAGTTCAGCAGTACg--------------------G-strand (upper strand)-------------------------
gatccgatatcACTAGTGTACACTGATCACATGTACTAGAATACCAGGTTACTATGAATGAGTACGATAGCTATCAGATGTACTACTACTAGATACACGTTGTCATTAGTCACTAGGAT
CAGTACTACTTCGATCACTAGATGTTACAGTCAGAGTCCTCTTAGATTGATACAGTATACTACATGACATATTAGTCCAGTACATACAGACATGATTCACGTGACATTATGAGTACC
ATCATTCAGATCAGTCACCATTACTGTGAGATACTAGCTAGAGTAGACTACTGATCGACTACATGTCAGTCTCAGTACGTTGTATCATAGTCACATTACATCATGTATGCTATTCAG
ATTGAGTTAACGGTACCGTCTcagcttGCACTGAGAGCGCGGCCTCTCAGTGCtgAGACGGTACCGTTAACTCAATCTGAATAGCATACATGATGTAATGTGACTATGATACAACGT
ACTGAGACTGACATGTAGTCGATCAGTAGTCTACTCTAGCTAGTATCTCACAGTAATGGTGACTGATCTGAATGATGGTACTCATAATGTCACGTGAATCATGTCTGTATGTACTG
GACTAATATGTCATGTAGTATACTGTATCAATCTAAGAGGACTCTGACTGTAACATCTAGTGATCGAAGTAGTACTGATCCTAGTGACTAATGACAACGTGTATCTAGTAGTAGTAC
ATCTGATAGCTATCGTACTCATTCATAGTAACCTGGTATTCTAGTACATGTGATCAGTGTACACTAGTgatatcg------------------------C-strand (lower strand)---------------------
aattcGTACTGCTGAACTATAGTTATCTAGTAGTCGATCAAGTAGACAATCATAGCATTAAGATGACATCAGAGTGTTACGAAGCTGACTACAGATAGTAGAAGATTCTCTCACTCA
AGTAGTCATGTCCTAGATACTGTACTGATCGATCTACTGATGATGCTGACTCAGAGTGTACTTGATAGTAACTGTAGTCATACATTCATGTGCTGATGTTAGACTGAGATACGTCA
GAGTGACTAGCTATGATAATCGCTATGTCAGTGTCTGTCAGGTTCATCTTGTTAGATGCTCGTACTTAGTAATACTGTCACTCTAGAATAGTAGCAGTCAGTCATAGCATGGTCATA
GTCAATCGATCTATCTACTTCAGTGTATACTTCTACTCCTGTCGTGTATGATCAGATCTGAGCTATGTATCTTCAGTACTCACTAGACTTATGAGGATAGTCTGACAGTACATCAAGA
TGGATCTGATCATGACATGAGTACTACAGTTCTAAGACTCTGTTACATATAGAGATGCTGATGGTACTGTCAGTCATTAGTATCTACTGTAGTCTTGACTCTAAGGCCTGAGCTCgT
ctccattcGAAGAGCACCAGAAAGACCAAAAGACACAAAGGGTCAGTGCTGCAACCCACTTCCTAATCTGTCATCTTCTG-3'

**Supplementary Data 2: Sequence of the common region of the DNA hairpin.** The hairpin single stranded loop sequence is shown in red. Sequences of the Y shape which allows tethering the hairpin to the surface and to the magnetic bead are shown in blue. Sequences belonging to the PS005 backbone are shown in brown. In black is the region where G4 sequence is inserted.

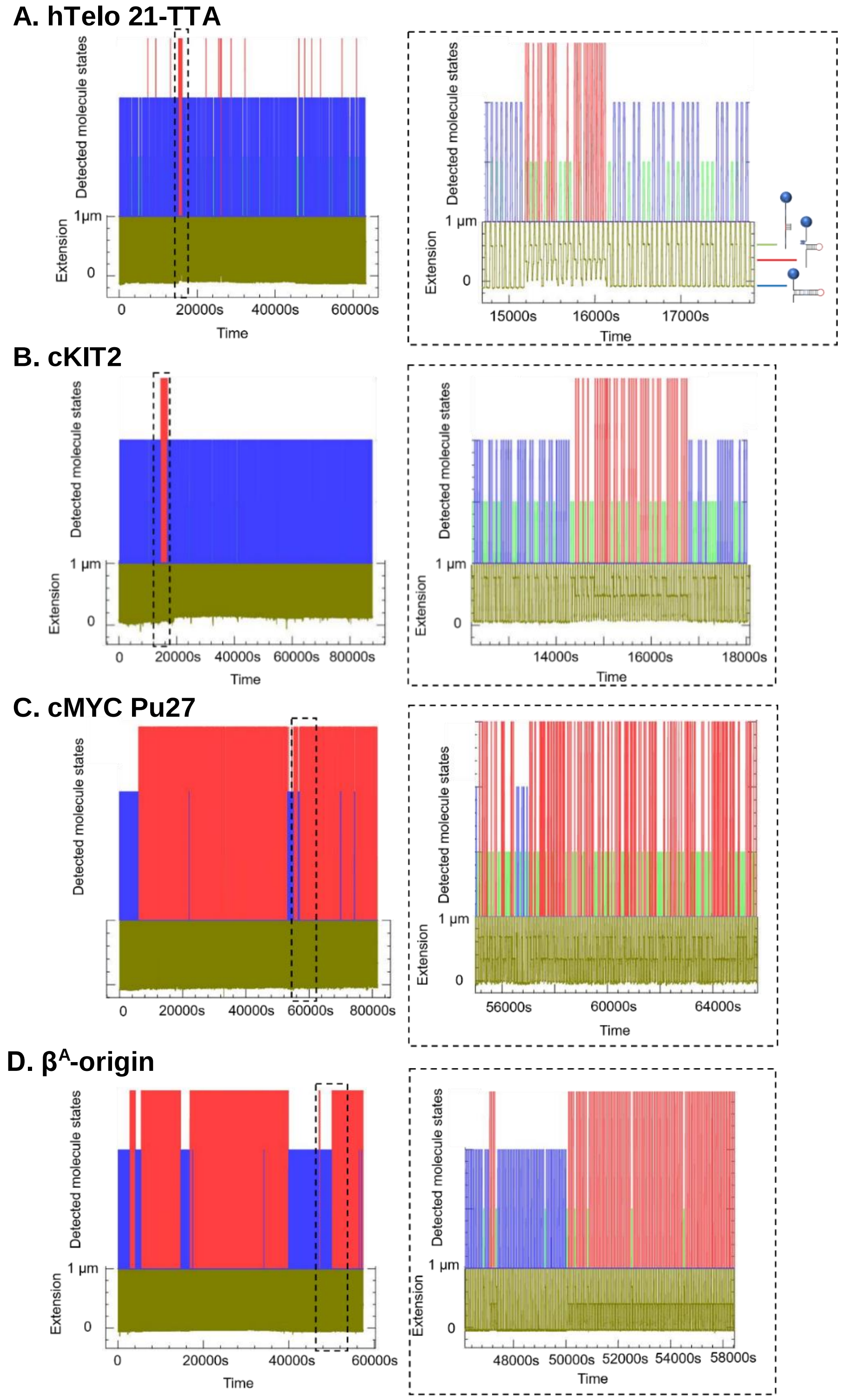


**Supplementary Data 3: Automatic detection of molecular states from full-time courses of recorded DNA extension traces obtained with hTelo 21-TTA (A), cKIT2 (B), cMYC Pu27 (C) and $\beta^A$ origin (D).** The left panel shows the full course of an experiment from one bead. Zoom on the dashed timeframe in the left panel is shown on the right. Traces of DNA extension are shown in khaki green. Blockages of the hairpin by a G4 structure or by the binding of oligonucleotide to its loop were automatically detected for each cycle. During hairpin refolding at 7 pN ($F_{hold}$), three detected molecule states are complete refolding of the hairpin (0 µm extension state is indicated in blue), binding of blocking oligonucleotide (0.8 µm extension state is in green) and G4 folding (0.4 µm extension state is indicated in red).

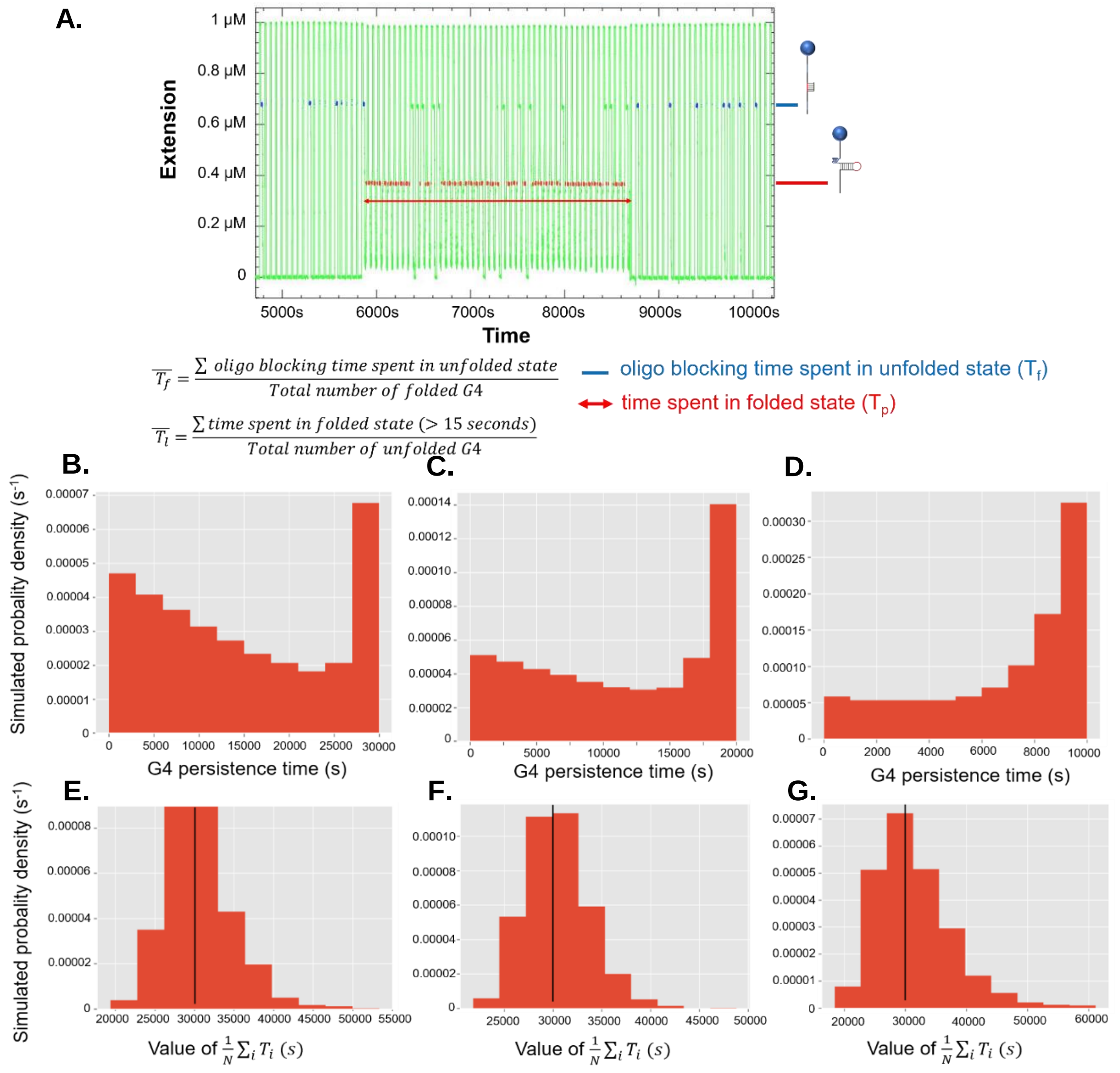


**Supplementary Data 4: Calculation of $\overline{T_f}$ and $\overline{T_p}$.**

**A. Diagram explaining the calculation method of the mean of G4 folding ($\overline{T_f}$) and G4 persistence times ($\overline{T_p}$). B-G. Simulations illustrating the impact of the experiment time of $\overline{T_f}$ and $\overline{T_p}$.** Experiments are simulated on 100 beads by taking single exponential distributions for the probabilities of folding ($\overline{T_f}$ = 100 s) /unfolding ($\overline{T_p}$ = 30000s) of the G-quadruplex as well as for the probabilities of binding ($\overline{T_{bound}}$ = 30 s)/unbinding ($\overline{T_{unbound}}$ = 10s) of the oligonucleotide.

The distribution of G4 lifetimes averaged over 1000 experiments are drawn for different measurement times in panel **B** ($T_{exp}$ = 10000s), **C** ($T_{exp}$ = 20000s) and **D** ($T_{exp}$ = 30000s). These figures show the impact of a short experiment time on the observed $T_p$ distributions, when the experiment time is smaller or comparable to the persistence time of a given G-quadruplex.

The panel **E, F** and **G** show the corresponding distribution over 1000 experiments of the resulting $\overline{T_p}$ as inferred on one experiment by using the method in the paper, that is by dividing the whole time where a G-quadruplex is observed by the number of unfolding events (described above in **A**). They validate the accuracy of the inference process even in cases when the $T_p$ is larger than the experiment time. Black lines represent the simulated $\overline{T_p}$ of the G4 (30000s), i.e. the correct value to be inferred. Standard deviation of the inferred value is respectively 3250s, 4310s and 6500s. The error of the inference increases as the measurement time decreases because of the corresponding decrease of the number of observed events.

**Supplementary Data 5: Error estimations of folding and persistence times**

The method used in this paper involves a periodic testing of the presence of a G-quadruplex in the substrate every cycle, each cycle lasting a time $T_c$. During a fraction of this cycle ($\alpha_i T_c$), the presence of the G4 cannot be detected, either because the hairpin is open or because the G4 is embedded in double-stranded DNA. Thus, there is a small probability that a G4 structure unfolds and then refolds during this time $\alpha_i T_c$. Then, we would observe two successive blockages at the G4 position that we would consider as a unique structure while in reality, two successive folding and unfolding events happened. In this section, we assess the error made on the estimation of the kinetic parameters of G4 folding ($k_f$) and unfolding ($k_u$) due to the possibility of these hidden events.

Given the knowledge that a G-quadruplex is present in the molecule at $t_0$, the probability $P_1$ that there is still a structure at a time $t_0 + \alpha_i T_c$, independently of whether it unfolds and refolds one or several times is:

$$P_1 = \frac{k_f}{k_u + k_f} + \frac{k_u}{k_u + k_f} e^{-(k_f+k_u)\alpha_i T_c}$$

Thus, the probability to detect a G4 during $N$ successive cycles is:

$$\begin{aligned} P(N) &= \left[\underbrace{\left(e^{-k_u(1-\alpha_i T_c)}\right)}_{P_0}\left(\underbrace{\frac{k_f}{k_u + k_f} + \frac{k_u}{k_u + k_f} e^{-(k_f+k_u)\alpha_i T_c}}_{P_1}\right)\right]^N \\ &= \left[\underbrace{\left(e^{-k_u(1-\alpha_i T_c)}\right)}_{P_0}\left(\underbrace{1 - (1 - e^{-(k_f+k_u)\alpha_i T_c}) \frac{k_u}{k_u + k_f}}_{P_1}\right)\right]^N, \end{aligned}$$

where $P_0$ is the probability that the G4 does not unfold during the part of the cycle where it is detectable and $P_1$ is defined just above. In particular, the probability to observe the G4 during N cycles if it were detectable at all times ($\alpha_i = 0$) would be $e^{-k_u N T_c}$.

Using the inequality $(1 + x)^N \leq e^{Nx}$, one gets:

$$P(N) < \left(e^{-k_u(1-\alpha_i T_c)}\right)^N e^{-N(1-e^{-(k_f+k_u)\alpha_i T_c})\frac{k_u}{k_u+k_f}}$$

Now considering that $e^{-x} < 1 - x + x^2/2$, we get:

$$\begin{aligned} P(N) &< \left(e^{-k_u(1-\alpha_i T_c)}\right)^N e^{-N\frac{k_u}{k_u+k_f}((k_f+k_u)\alpha_i T_c + (k_f+k_u)^2 \alpha_i{}^2 T_c{}^2)} \\ &< e^{-k_u T_c N} e^{-N(k_f+k_u)k_u \alpha_i^2 T_c^2/2} \\ &< e^{-T_c N \underbrace{k_u(1-(k_f+k_u)\alpha_i^2 T_c/2)}_{k_{u,obs}}} \end{aligned}$$

This means that the observed folding kinetics $k_{u,obs}$ is slightly different from the real rate $k_{u,real}$, with:

$$k_{u,real} > k_{u,obs} > k_{u,real}(1 - (k_f + k_u)\alpha_i^2 T_c/2)$$

In particular, this means that we slightly overestimate the persistence time $T_p = \frac{1}{k_u}$ and that the amplitude of the overestimation verifies:

$$T_{p,real} < T_{p,obs} < \frac{T_{p,real}}{1 - (k_f + k_u)\alpha_i^2 T_c/2}$$

Conversely, we make a similar error on $T_f = \frac{1}{k_f}$, as it is possible that between two successive detections steps, one was folded and then unfolded:

We thus have,

$$T_{f,real} < T_{f,obs} < \frac{T_{f,real}}{1-(k_f+k_u)\alpha_i^2 T_c/2}$$

For our data, these corrections are negligible (< 10%), except for the $\beta^A$-origin sequence that folds very rapidly. This overestimation of the measured times is taken into account in all our error bars. We add to the low error bars of our data the maximum relative error that can be made due to these hidden events: $(k_f + k_u)\alpha_i^2 T_c^2/2$.

In all the data shown in this paper, we take as a lower error the sum of the probabilistic error due to the above effect and of the sampling errors (obtained through bootstrap-resampling of the data). However, we take as upper error the bootstrapped error only as the hidden events cannot induce any underestimation of the times:

$$\sigma_{up} = \sigma_{bootstrap}$$

$$\sigma_{down}^2 = \sigma_{bootstrap}^2 + max((k_{real} - k_{obs}))^2$$

A.

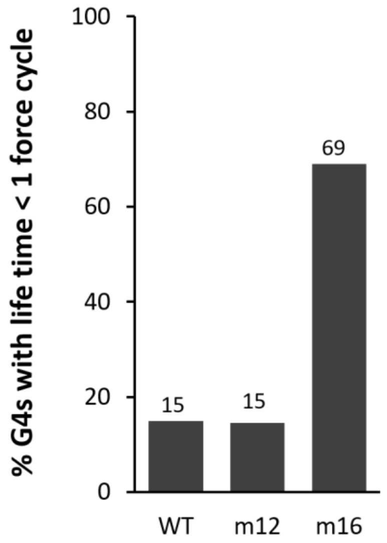


B.

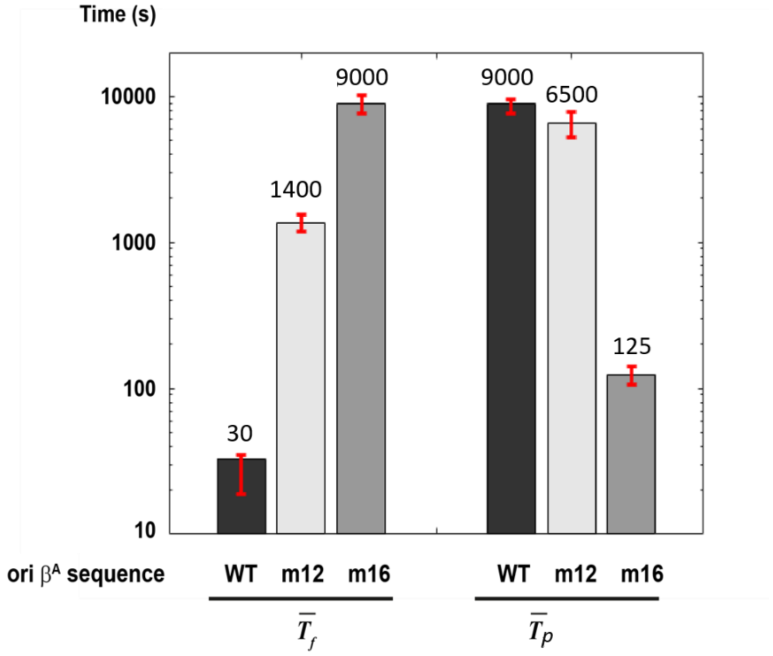


**Supplementary Data 6: Folding and persistence times of G4 structures formed by ori-β$^A$ wild type sequence and point mutants m12 and m16. A.** Percentage of G4 with persistence time less than 1 force cycle. **B**. Mean folding ($\overline{T_f}$) and persistence ($\overline{T_p}$) times of wild type (WT) and mutant ori-β$^A$ G4 forming sequences.

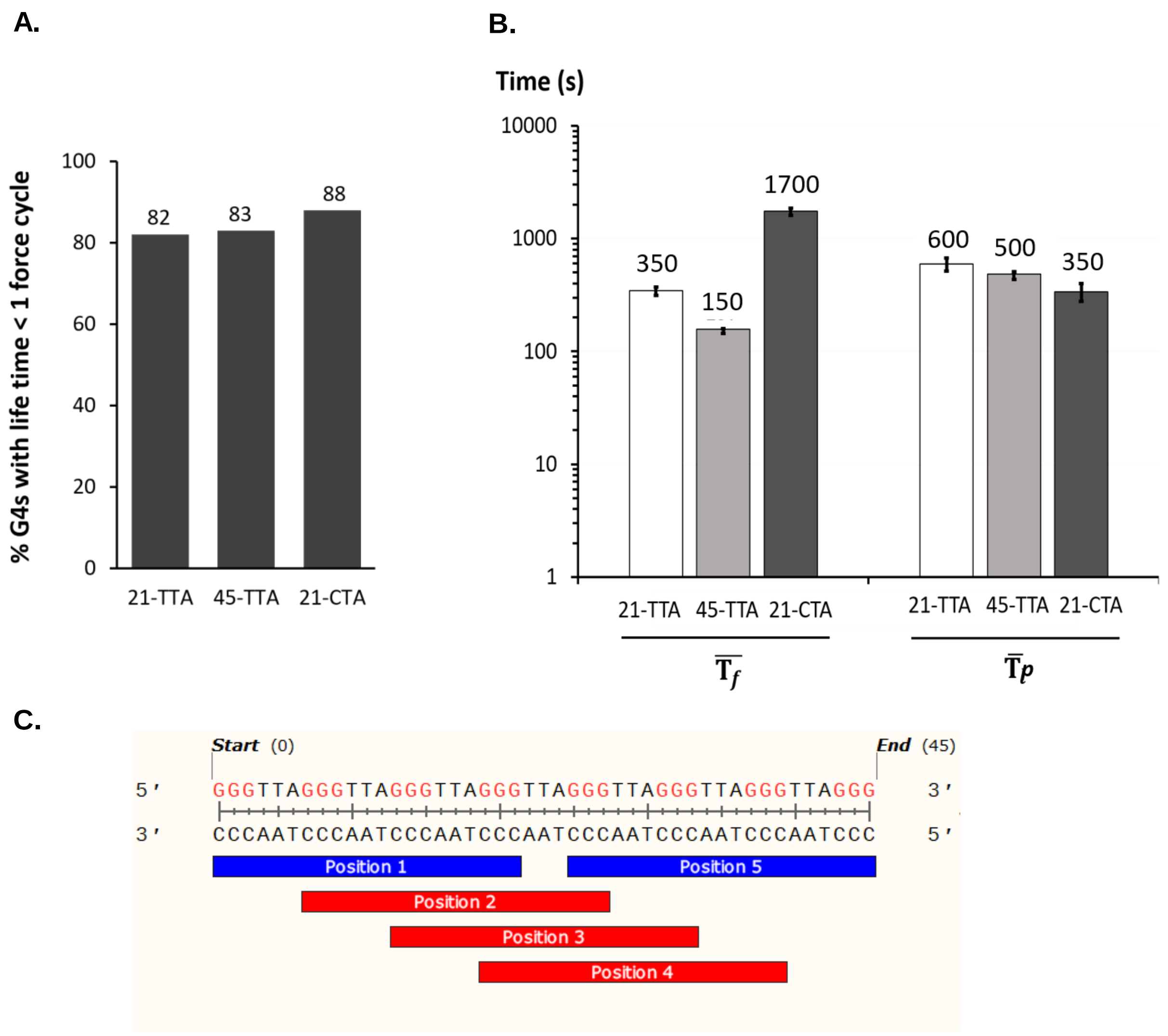


**Supplementary Data 7: Folding and persistence of G4 structures formed by human telomeric sequences. A.** Percentage of G4 with persistence time less than 1 force cycle. **B**. Mean folding ($\overline{T_f}$) and persistence ($\overline{T_p}$) times of human telomeric sequences. **C.** Different possible G4 nucleation sites folded by 45-TTA sequence.

**A.**

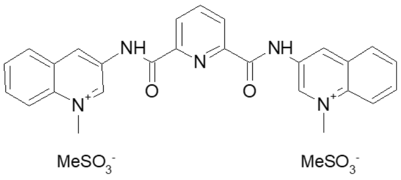


**360B**

**B.**

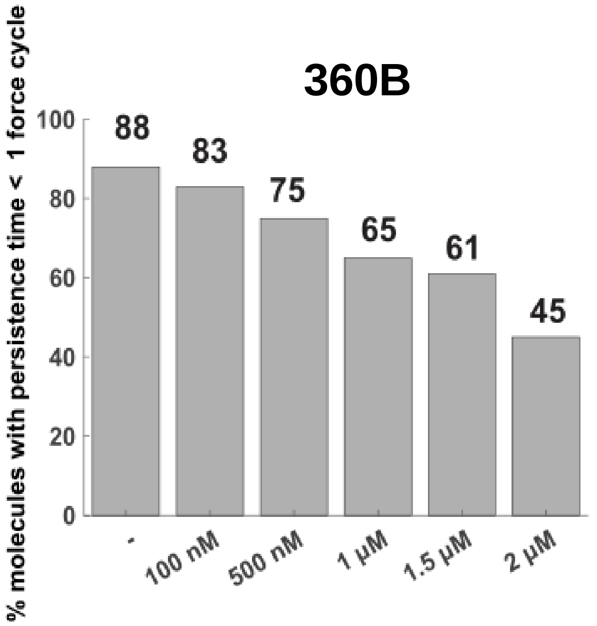


**C.**

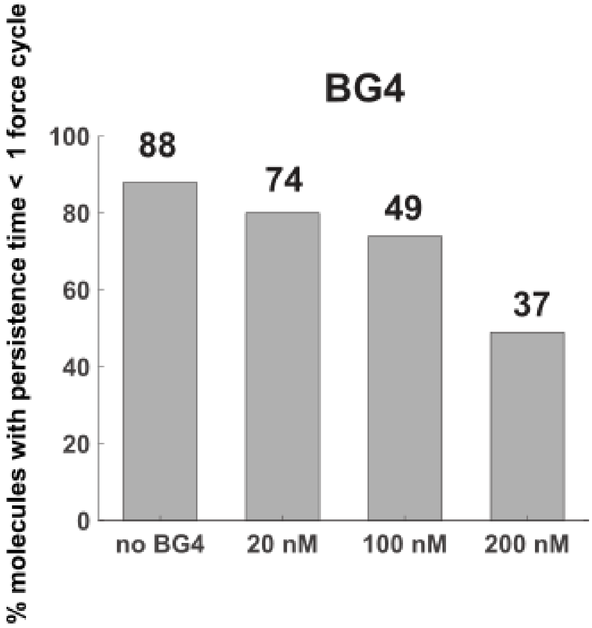


**Supplementary Data 8: Effect of G4 ligand 360B and BG4 antibody on the persistence of 21-CTA G4.** (**A**) Chemical formula of 360B ligand. Percentage of 21-CTA G4 with persistence time less than 1 force cycle in presence of increasing concentration of 360B (**B**) and BG4 antibody (**C**).

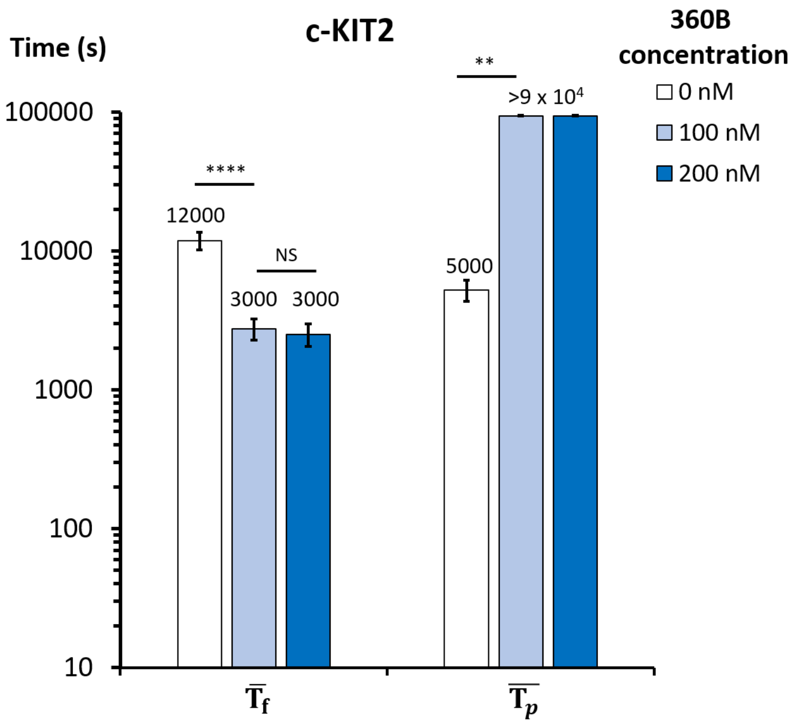


**Supplementary Data 9: The mean of folding and persistence time of c-KIT2 at different concentrations of 360B.** P-values are from ~ 100 events and are calculated using two-tailed t-test. NS stands for non-significative ($p > 0.1$). *** $p \leq 0.005$ and **** $p < 0.00001$

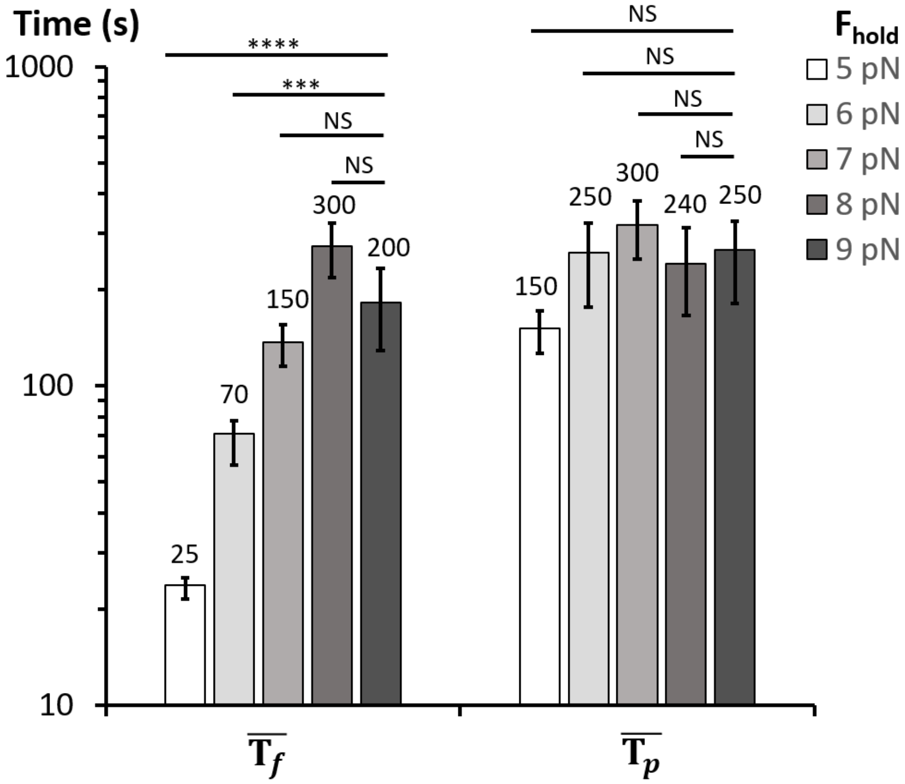


**Supplementary Data 10: Mean folding time and persistence time of human telomeric G4 forming sequences (45-TTA) at different holding forces.** We performed different cycle experiments on human telomeric 45-TTA sequence by varying the $F_{hold}$ from 5 to 9 pN (force that we can observe blocking oligonucleotide and G4 formation). The results show that the lifetime of 45-TTA G4 did not depend on the applied $F_{hold}$. However, its folding time increased with the force until reaching a plateau at ~ 7-9 pN. We have then chosen the holding force ($F_{hold}$) at 7 pN for all G4 experiments where we can have reasonable oligo-blocking time (about 10-15s) and where most of DNA hairpins were closed in the absence of oligonucleotide and G4 formation. P-values are from ~ 100 events and are calculated using two-tailed t-test. NS stands for non-significative (p > 0.1). ******* p ≤ 0.0001 and **** p ≤ 0.00001

| Name | Description | Sequence (5’ □ 3’) |
|---|---|---|
| PS124 | Hairpin loop | GCTTGCACTGAGAGCGCGGCCTCTCAGTGC |
| Treble | Tri-biotinylated 5’ end oligonucleotide allowing binding of the hairpin to a streptavidin covered magnetic bead | *Biotin(x3)*_ATGGTCAAGCATGCCGCTTTTCGGTTC CCGTGTGTCTTTTGGTCTTTCTGGTGCTCTTC |
| PS866 | 3’-end oligonucleotide allowing annealing of the hairpin to oligonucleotide PS867 | ATTCGAAGAGCACCAGAAAGACCAAAAGACACAA AGGGTCAGTGCTGCAACCCACTTCCTAATCTGTC ATCTTCTG |
| PS867 | Bridge oligonucleotide to link PS866 with PS625 | GTGTCTTTTGGTCTTTCTGGTGCTCTTCGAATCA GAAGATGACAGATTAGGAAGTGG |
| PS625 | sequence containing a 3’ DBCO group which covalently binds to the azide coated cover slip | ATTCGAAGAGCACCAGAAAGACCAAAAGACACA GACAGATATCGCGCTTCCTCCTACTTTGAATGCT AT_*DBCO* |
| Blocking oligonucleotide | 7 bp blocking oligonucleotide | GCCGCGC |

**Supplementary Table 1: Sequences of DNA oligonucleotides used for hairpin synthesis and for tethering hairpins to the azide coated coverslip.** In black are single-stranded DNA regions. The regions with identical colors (*i.e.* red, purple, orange and green) are complementary sequences.

| Name | Description | Sequence (5’ □ 3’) |
|---|---|---|
| hTelo 21-CTA | variant human telomeric sequence (single G4 unit) | AATTCTCATAGCATGATACCAATGGGCTAGGGCTAGGGCTAGGGATTGGTACGTAGACCATG |
| hTelo 21-CTA mut | mutated hTelo 21-CTA | AATTCTCATAGCATGATACCAATGcGCTAGcGCTAGcGCTAGcGATTGGTACGTAGACCATG |
| cMYC Pu27 | promoter of oncogene c-MYC | ATTCTCATAGCATGATAAGGGGAGGGTGGGGAGGGTGGGGAAGGTTGGTACGTAGACCATG |
| cmyc Pu27 mut | mutated c-MYC Pu27 | AATTCTCATAGCATGATAAGGGGAGcGTGccGAGcGTGccGAAGGTTGGTACGTAGACCATG |
| cKIT2 | promoter of oncogene c-KIT | ATTCTCATAGCATGATACCAATGGGCGGGCGCGAGGGAGGGGAATTGGTACGTAGACCATG |
| ckit2 mut | mutated c-KIT2 | AATTCTCATAGCATGATACCAATGcGCGcGCGCGAGcGAGccGAATTGGTACGTAGACCATG |
| $\beta^A$ ori | Chicken origin of replication $\beta^A$ | AATTCTCATAGCATGATACCAATGGGGGGGGGGGGGCGGGATGCATTGGTACGTAGACCATG |
| $\beta^A$ ori - M12 | mutated ori $\beta^A$ ($G_{12}$□A) | ATTCTCATAGCATGATACCAATGGGGGGGGGGGaGCGGGATGCATTGGTACGTAGACCATG |
| $\beta^A$ ori - M16 | mutated ori $\beta^A$ ($G_{16}$□A) | AATTCTCATAGCATGATACCAATGGGGGGGGGGGGGCGaGATGCATTGGTACGTAGACCATG |
| hTelo 21-TTA | human telomeric sequence (single G4 unit) | AATTCTCATAGCATGATACCAATGGGTTAGGGTTAGGGTTAGGGATTGGTACGTAGACCATG |
| hTelo 21-TTA mut | mutated hTelo 21-TTA | AATTCTCATAGCATGATACCAATGcGTTAGcGTTAGcGTTAGcGATTGGTACGTAGACCATG |
| hTelo 45-TTA | human telomeric sequence (two G4 units) | AATTCTCATGGGTTAGGGTTAGGGTTAGGGttaGGGTTAGGGTTAGGGTTAGGGAGACCATG |
| hTelo 45-TTA | mutated hTelo 45-TTA | AATTCTCATGcGTTAGcGTTAGcGTTAGcGttaGcGTTAGcGTTAGcGTTAGcGAGACCATG |

**Supplementary Table 2: G4 forming sequences and control sequences used in this study:** in red, guanines involved in the G4 structure, and in lower-cases, base mutated to destabilize G4 structures.
